# Comparative genomic analyses of *Lactococcus garvieae* isolated from bovine mastitis in China

**DOI:** 10.1101/2022.08.01.502421

**Authors:** Yushan Lin, Jinge Han, Herman W. Barkema, Yue Wang, Jian Gao, John P. Kastelic, Bo Han, Shunyi Qin, Zhaoju Deng

**Affiliations:** Department of Clinical Veterinary Medicine, College of Veterinary Medicine, China Agricultural University, Beijing, 100193, P. R. China; College of Animal Science and Veterinary Medicine, Tianjin Agricultural University, Tianjin, 300384, P. R. China; Department of Production Animal Health, Faculty of Veterinary Medicine, University of Calgary, Calgary, AB, Canada, T2N 4N1

**Keywords:** bovine mastitis, *Lactococcus garvieae*, population structure, virulence genes, antimicrobial resistance, host adaptation

## Abstract

*Lactococcus garvieae* is an emerging zoonotic pathogen, but there are few reports regarding bovine mastitis. The recent prevalence of *L. garvieae* poses an increasing disease threat and global public health risk. A total of 39 *L. garvieae* isolates were obtained from 2899 bovine clinical mastitis milk samples in 6 provinces of China from 2017 to 2021. Five clonal complexes were determined from 32 MLST types of *L. garvieae*; ST46 was the predominant sequence type and 13 novel MLST types were identified. All isolates were resistant to chloramphenicol and clindamycin, but susceptible to penicillin, ampicillin, amoxicillin-clavulanic acid, imipenem, ceftiofur, enrofloxacin, and marbofloxacin. Based on genomic analyses, *L. garvieae* had 6310 genes, including 1015, 3641 and 1654 core, accessory and unique genes. All isolates had virulence genes coding for collagenase, fibronectin-binding protein, Glyceraldehyde-3-phosphate dehydrogenase, superoxide dismutase and NADH oxidase. Most isolates had *lsaD* and *mdtA* AMR genes. Based on COG results, the functions of defense, transcription and replication, recombination and repair were enhanced in unique genes, whereas functions of translation, ribosomal structure and biogenesis were enhanced in core genes. The KEGG functional categories enriched in unique genes included human disease and membrane transport, whereas COG functional categories enriched in core genes included energy metabolism, nucleotide metabolism and translation. No gene was significantly associated with host specificity. In addition, core genome SNPs analysis suggested potential host adaptation of some isolates in several sequence types. Therefore, this study characterized *L. garvieae* isolated from mastitis and assessed host adaptation of *L. garvieae* to various hosts.

**IMPORTANCE:** This study provides important insights on bovine mastitis key topic pathogen *Lactococcus garvieae*, which constitutes mastitis concerns. However, comprehensive genomic analyses of *L. garvieae* from dairy farms have not been performed. This study gives a detailed and comprehensive novel feature in *L. garvieae*, an important but poorly characterized bacterium, recovered in the past 5 years in 6 Chinese provinces. We documented diverse contributory genetic processes, including predominant sequence type ST46 and 13 novel MLST types. *L. garvieae* had 6310 genes, including 1015, 3641 and 1654 core, accessory and unique genes. All isolates had virulence genes coding for collagenase, fibronectin-binding protein, Glyceraldehyde-3-phosphate dehydrogenase, superoxide dismutase and NADH oxidase, and resistant to chloramphenicol and clindamycin. Most isolates had *lsaD* and *mdtA* antimicrobial resistance genes. No gene was significantly associated with host specificity. This is the first absolute quantification of *L. garvieae* isolated from mastitis and identified host adaptation of *L. garvieae* to various hosts.

## INTRODUCTION

Bovine mastitis is a prevalent and costly disease on dairy farms worldwide (**1, 2**). It is a multifactorial disease, often caused by bacteria (**2**). Bacterial pathogens associated with bovine mastitis are broadly classified as major (*Staphylococcus aureus*, *Escherichia coli*, *Streptococcus agalactiae, Streptococcus dysgalactiae, Streptococcus uberis*, *Enterococcus* spp. etc.) and minor pathogens (Non*-aureus Staphylococci* spp., *Lactococcus* spp., Corynebacterium spp., etc.) (**3**). Most studies have focused on major pathogens, with only limited studies of minor pathogens.

*Lactococcus garvieae* is a zoonotic pathogen reported to cause infections in fish (**4**) and humans (**5–8**). It is also considered a minor pathogen for bovine mastitis, with transmission attributed to environmental reservoirs (**2**). There are limited reports on bovine mastitis caused by *L. garvieae* (**9–15**), primarily descriptive studies of phenotypes or genotypes. However, detailed whole genome characterization of *L. garvieae* associated with bovine mastitis is lacking.

Predominant strain types of mastitis pathogens have been described for various pathogens (**16–20**). Elucidating population structure and diversity of mastitis pathogens informs evidence-based mastitis control programs that target those prevalent strain types.

Bacterial pathogenicity is primarily determined by virulence factors; some facilitate adhesion and invasion, whereas antimicrobial resistance, particularly multi-drug resistance, is an important threat to public health (**21**). For *L. garvieae,* several virulence factors and antimicrobial resistance genes were identified using traditional methods (e.g., PCR) targeted at specific virulence genes (**22–24**). However, a comprehensive profiling of its virulence and antimicrobial resistance genes are lacking.

Host adaptation of bovine mastitis associated pathogens have been reported for *Staphylococcus aureus* (**25**) and *Streptococcus agalactiae* (**26**). Infections caused by *L. garvieae* in humans, fish and cattle have been reported, but potential host adaptation of *L. garvieae* has not been studied. Therefore, our objectives were to: 1) resolve the population structure; 2) identify virulence genes and antimicrobial resistance genes; and 3) determine genes associated with host specificity.

## RESULTS

A total of 39 *L. garvieae* isolates from 2899 clinical mastitis composite milk samples collected from 13 large dairy farms in 6 provinces in Northern China from April 2017 to September 2021. Detailed information of these isolates is provided in **Table 1**.

**Table 1.** *Lactococcus garvieae* isolates (n=39) recovered from bovine mastitis from 2899 bovine mastitis (CM) milk samples collected from farms in China.

| Isolate | Province | Farm | Date |
| --- | --- | --- | --- |
| Shanxi-A-01 | Shanxi | A | 2018.1.29 |
| Hebei-B-02 | Hebei | B | 2018.4.9 |
| Hebei-C-03 | Hebei | C | 2018.5.11 |
| InnerMongolia-D-04 | Inner Mongolia | D | 2018.6.26 |
| Hebei-E-05 | Hebei | E | 2018.7.16 |
| InnerMongolia-F-06 | Inner Mongolia | F | 2019.6.25 |
| InnerMongolia-F-07 | Inner Mongolia | F | 2019.6.25 |
| Hebei-G-08 | Hebei | G | 2019.6.29 |
| Hebei-G-09 | Hebei | G | 2019.6.29 |
| Hebei-G-10 | Hebei | G | 2019.6.29 |
| Gansu-H-11 | Gansu | H | 2020.8.12 |
| Gansu-H-12 | Gansu | H | 2020.8.12 |
| Ningxia-I-13 | Ningxia | I | 2020.8.12 |
| Ningxia-I-14 | Ningxia | I | 2020.8.12 |
| Ningxia-I-15 | Ningxia | I | 2020.8.12 |
| Ningxia-I-16 | Ningxia | I | 2020.8.12 |
| Ningxia-I-17 | Ningxia | I | 2020.8.24 |
| Ningxia-I-18 | Ningxia | I | 2020.8.24 |
| Ningxia-I-19 | Ningxia | I | 2020.8.24 |
| Ningxia-I-20 | Ningxia | I | 2020.8.24 |
| Hebei-B-21 | Hebei | B | 2020.8.27 |
| Hebei-B-22 | Hebei | B | 2020.8.27 |
| Ningxia-I-23 | Ningxia | I | 2020.10.2 |
| Shanxi-J-24 | Shanxi | J | 2020.11.1 |
| Shanxi-J-25 | Shanxi | J | 2020.11.1 |
| Ningxia-I-26 | Ningxia | I | 2020.11.6 |
| Ningxia-I-27 | Ningxia | I | 2020.11.6 |
| Ningxia-I-28 | Ningxia | I | 2021.1.29 |
| Shandong-K-29 | Shandong | K | 2021.6.9 |
| Shandong-K-30 | Shandong | K | 2021.6.9 |
| Hebei-L-31 | Hebei | L | 2021.6.10 |
| Hebei-L-32 | Hebei | L | 2021.6.10 |
| Hebei-L-33 | Hebei | L | 2021.6.10 |
| Hebei-L-34 | Hebei | L | 2021.6.10 |
| Hebei-L-35 | Hebei | L | 2021.6.10 |
| Shandong-M-36 | Shandong | M | 2021.6.28 |
| Shandong-M-37 | Shandong | M | 2021.6.28 |
| Hebei-B-38 | Hebei | B | 2021.7.12 |

### MLST and Minimum Spanning Tree

MLST analysis assigned the 86 *L. garvieae* isolates into 32 STs (**Table 2**), of which 13 were novel STs: ST46 to ST58. The most common sequence type was ST46 (n = 13), followed by ST48 (n = 9). The 32 STs were grouped into 5 CCs and 18 singletons: ST 3, 4, 13 and 38 into CC 1; ST 16 and 17 into CC 2; ST 10, 12, 21 and 35 into CC 3; ST 49 and 52 into CC4; and ST 47 and 50 into CC 5 (**Fig. 1**). Among the 39 isolates, 6 and 9 isolates formed clonal complex CC4 and CC5, respectively, whereas, the remaining 24 isolates were singletons.

**FIG 1.**
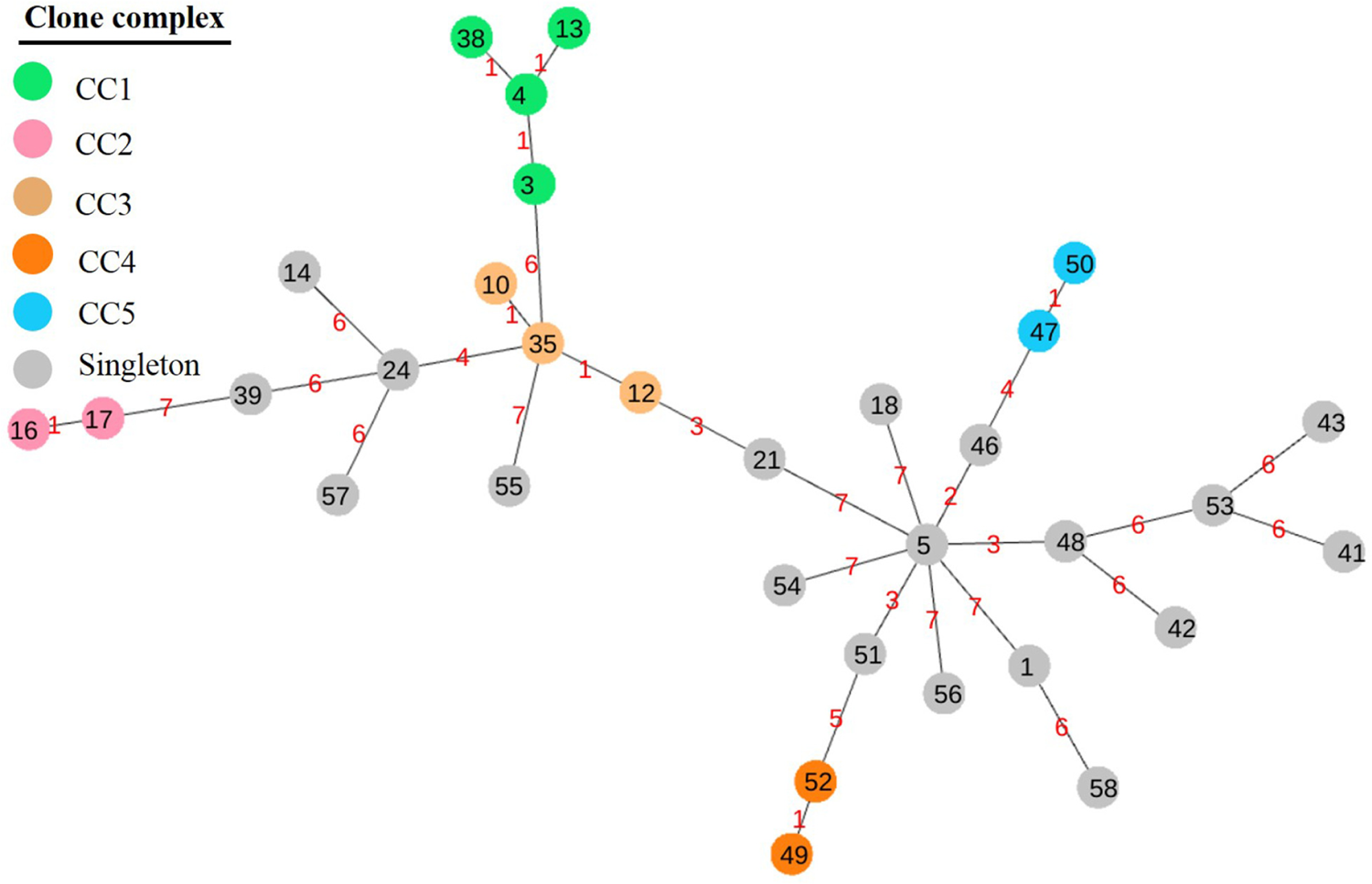
Minimum spanning tree based on Multi-Locus Sequence Typing for 86 *Lactococcus garvieae* isolates involving 32 sequence types performed by geoBURST algorithm and visualized by PhyloViz. Five clonal complexes (CC1-CC5) were clustered with similar STs (5 - 7 shared alleles); the number between nodes indicates the number of distinct alleles within them.

**Table 2.**
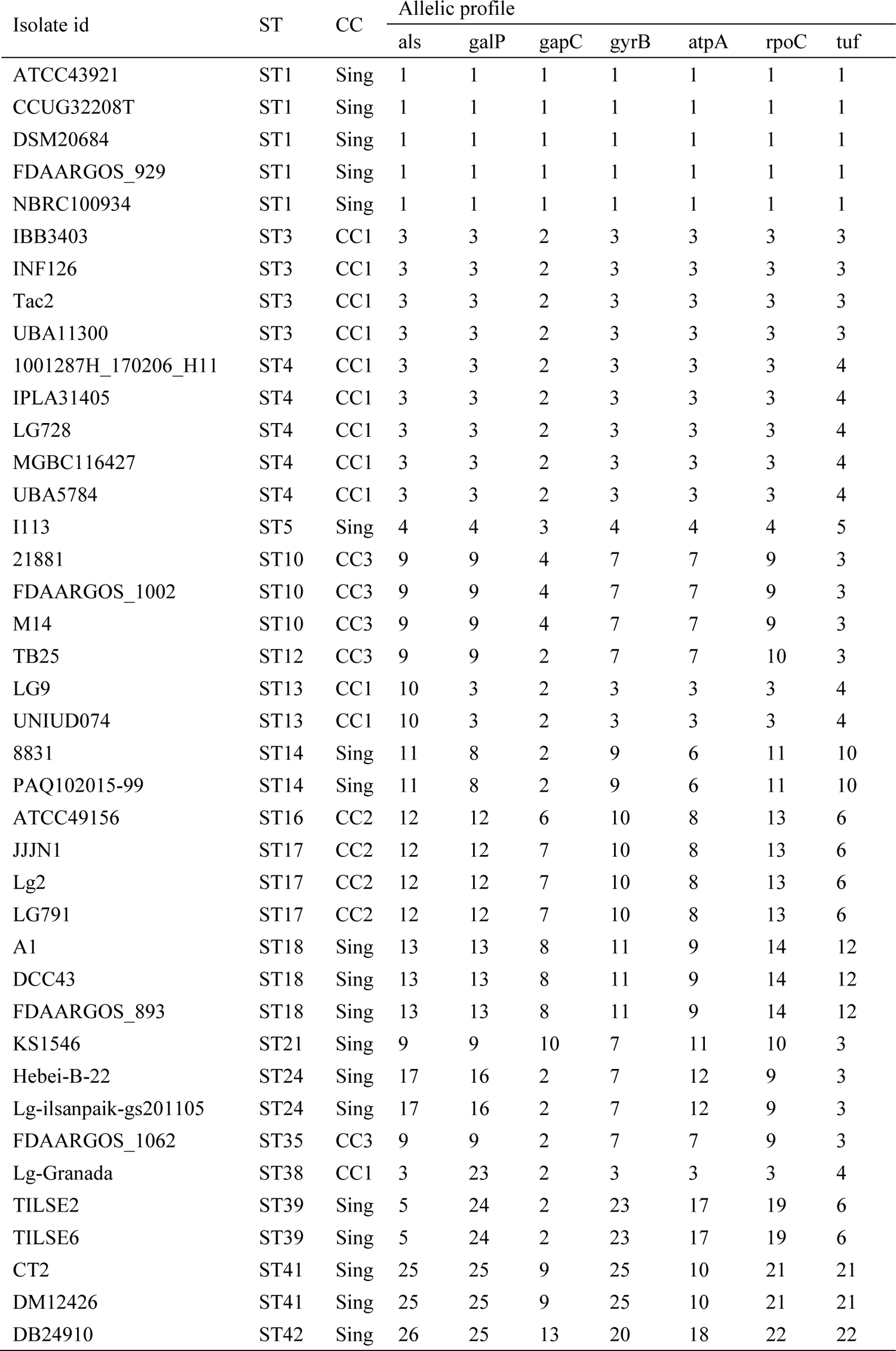
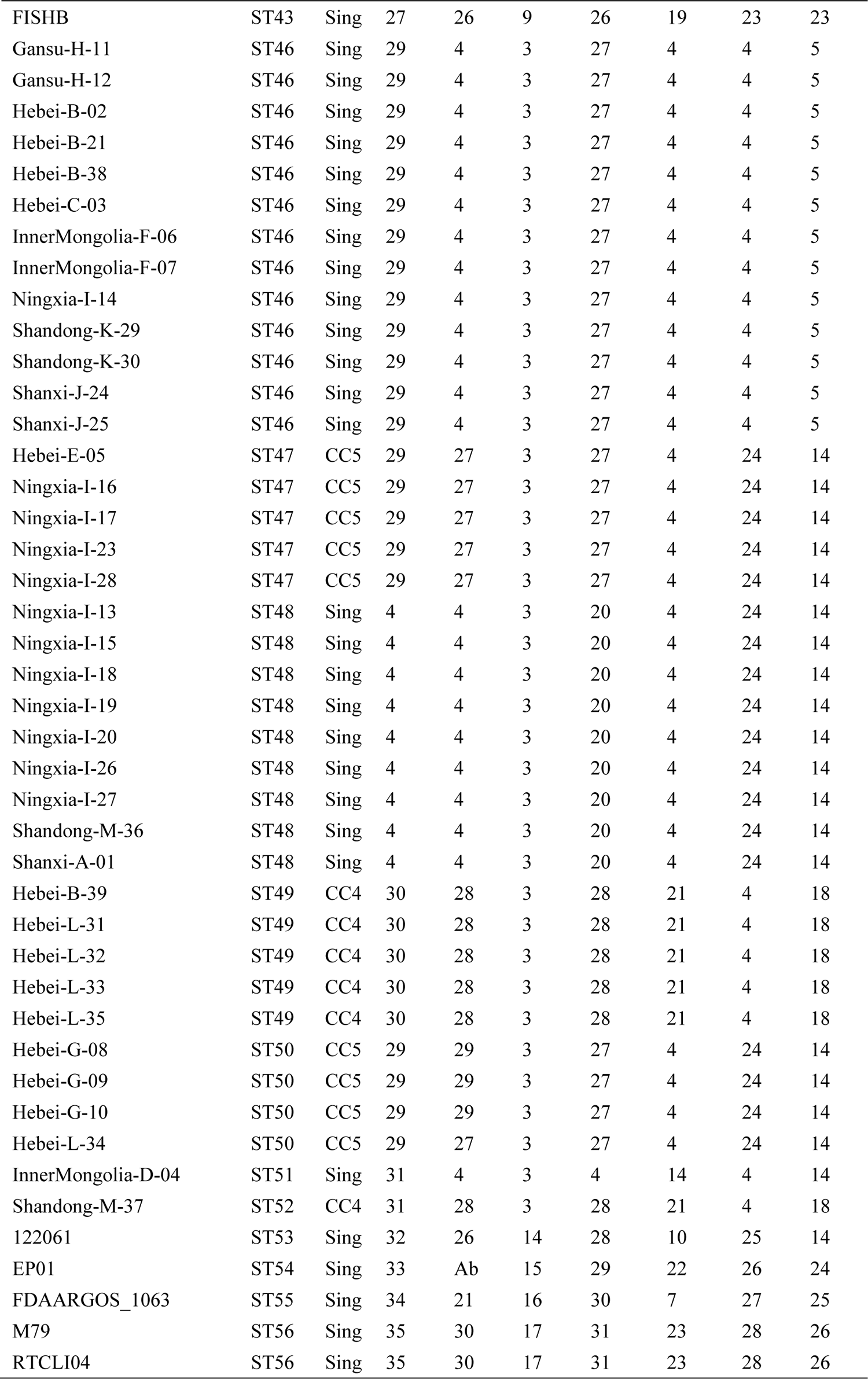
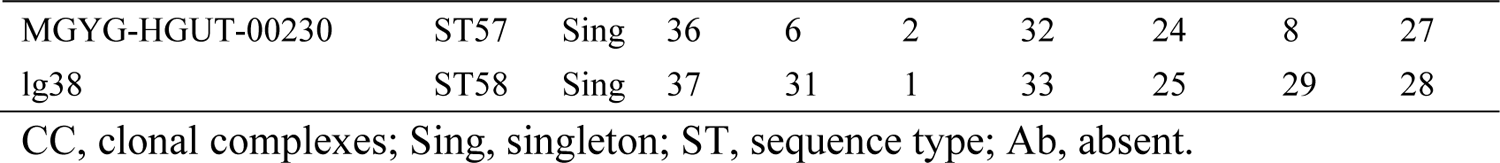
Allelic profiles, sequence types and clonal complexes of 86 *Lactococcus garvieae* isolates.

### Antimicrobial Resistance Profile and Genes

Antimicrobial resistance patterns of *L. garvieae* are listed in **Table 3**. All isolates were resistant to chloramphenicol and clindamycin. There were also high resistance rates for amikacin (90%), cefpodoxime (82.5%), cefazolin (45%), and gentamicin (37.5%), whereas few isolates (5%) were resistant to erythromycin. Meanwhile, all isolates were susceptible to penicillin, ampicillin, amoxicillin-clavulanic acid, imipenem, ceftiofur, enrofloxacin and marbofloxacin. Regarding multidrug resistance, there were 5, 14, 12, 6, 2 and 1 isolates resistant to 3, 4, 5, 6, 7, and 8 antimicrobials, respectively.

**Table 3.**
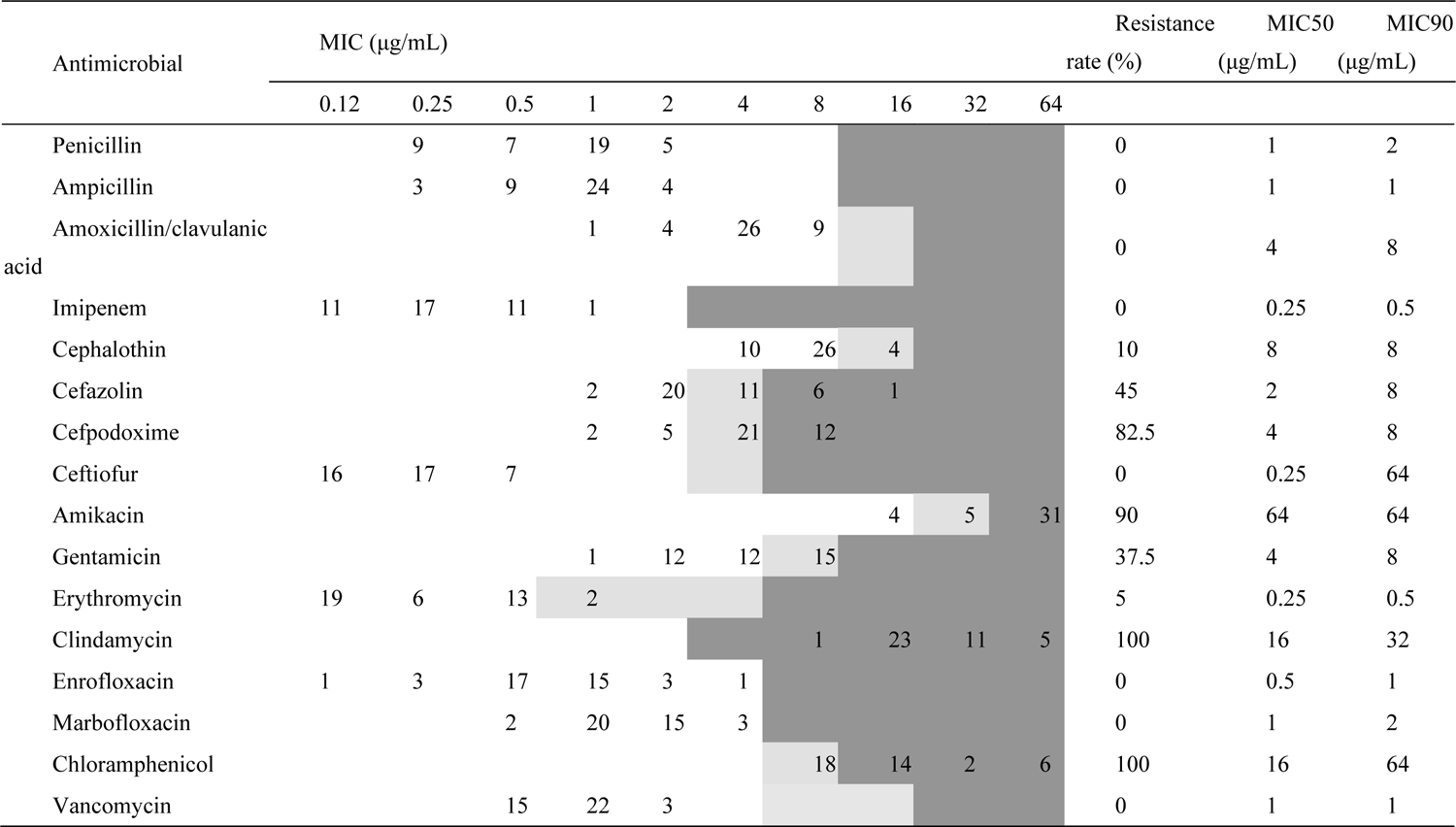
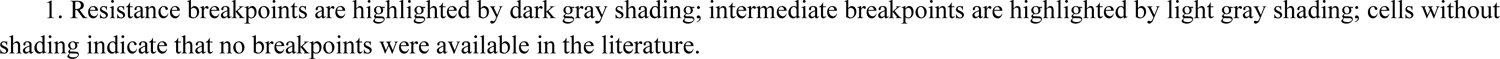
Minimum inhibitory concentration (MIC) of the 15 antimicrobials tested for 39 *Lactococcus garvieae* isolates isolated from clinical mastitis composite milk samples in China and control strain ATCC 43921 ^1^.

Distribution of antimicrobial resistance (AMR) genes with country, host and clonal complex information are presented in **Fig. 2**. The most common genotypic AMR markers were: (i) multidrug protein first reported in *L. garvieae*, represented by the *lsaD* gene (97% of isolates); (ii) resistance-nodulation-cell division antibiotic efflux pump, represented by the mdtA gene (97% isolates); and (iii) tetracycline-resistant ribosomal protection protein, coded by *tetS* gene (14 isolates, including 12 from current bovine samples). Other tetracycline-resistant genes*, tetL and tetM,* were identified in only 3 isolates. Three isolates, namely LG728, LG791 and MGYG-HGUT-00230, harbored the most abundant AMR genes, the gene number of which was 14, 7, 5 respectively. Genes *cat, dfrG, ermA* and *lunD* were only present in LG728, and gene *lnuD* was only in MGYG-HGUT-00230. Genes *ermB, fexA* and *optrA* were present in LG728 and LG791. Genes *acc(6’)-aaph(2’’)* and *ant*(*6*)*-la* were identified in LG728 and MGYG-HGUT-00230. However, no AMR genes were detected in 3 isolates that belonged to ST18: A1, DCC43 and FDAARGOS_893.

**FIG 2.**
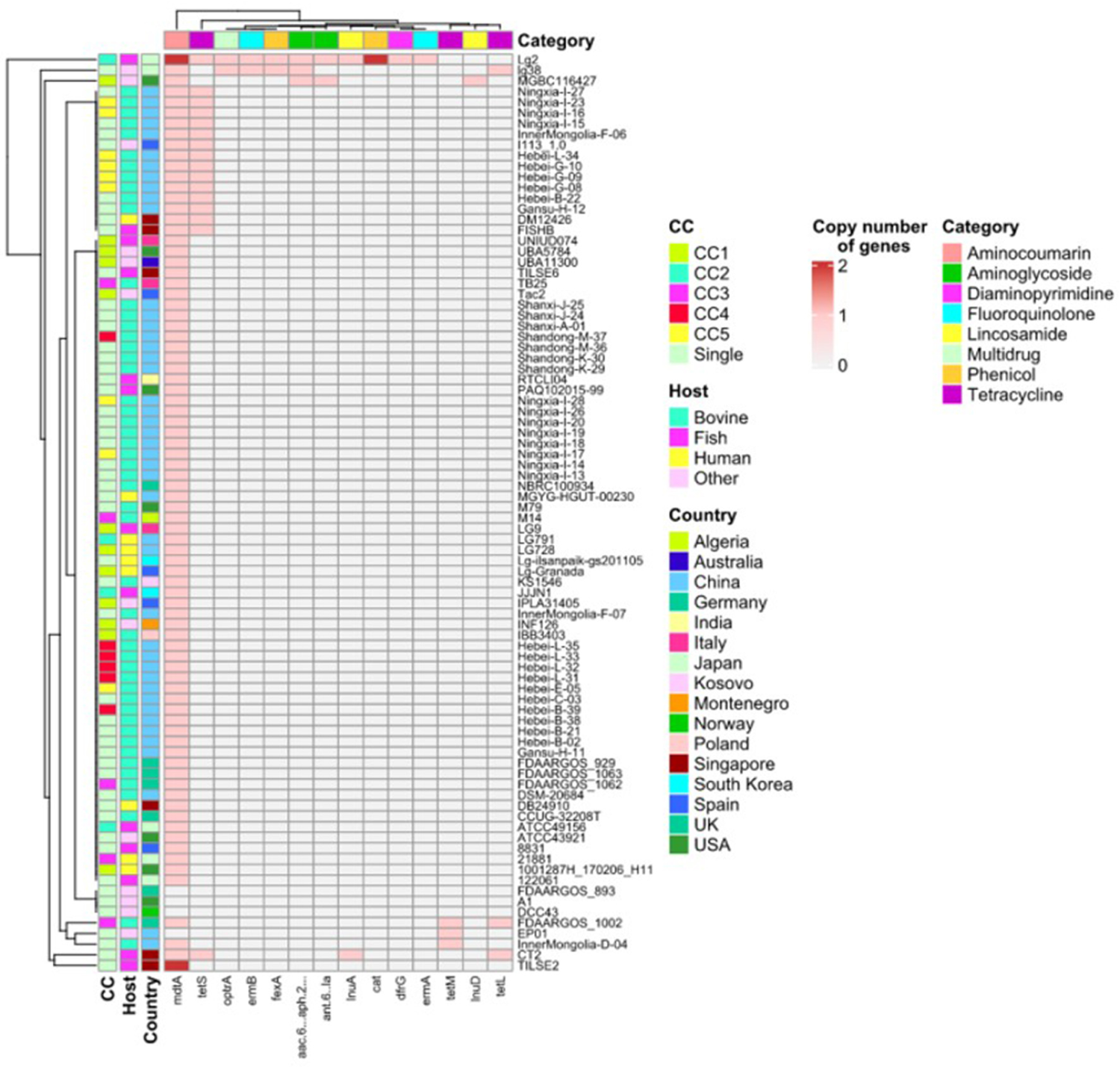
Distribution of antimicrobial resistance genes against each category of antimicrobial resistance, together with source of host, country, and clonal complex (CC) of 86 *Lactococcus garvieae* isolates.

### Virulence Genes

The occurrence and distribution of putative virulence genes are shown in **Fig. 3**. The putative virulence factors were classified into 5 functional categories: toxin, iron uptake, capsule formation, adherence, and enzyme.

**FIG 3.**
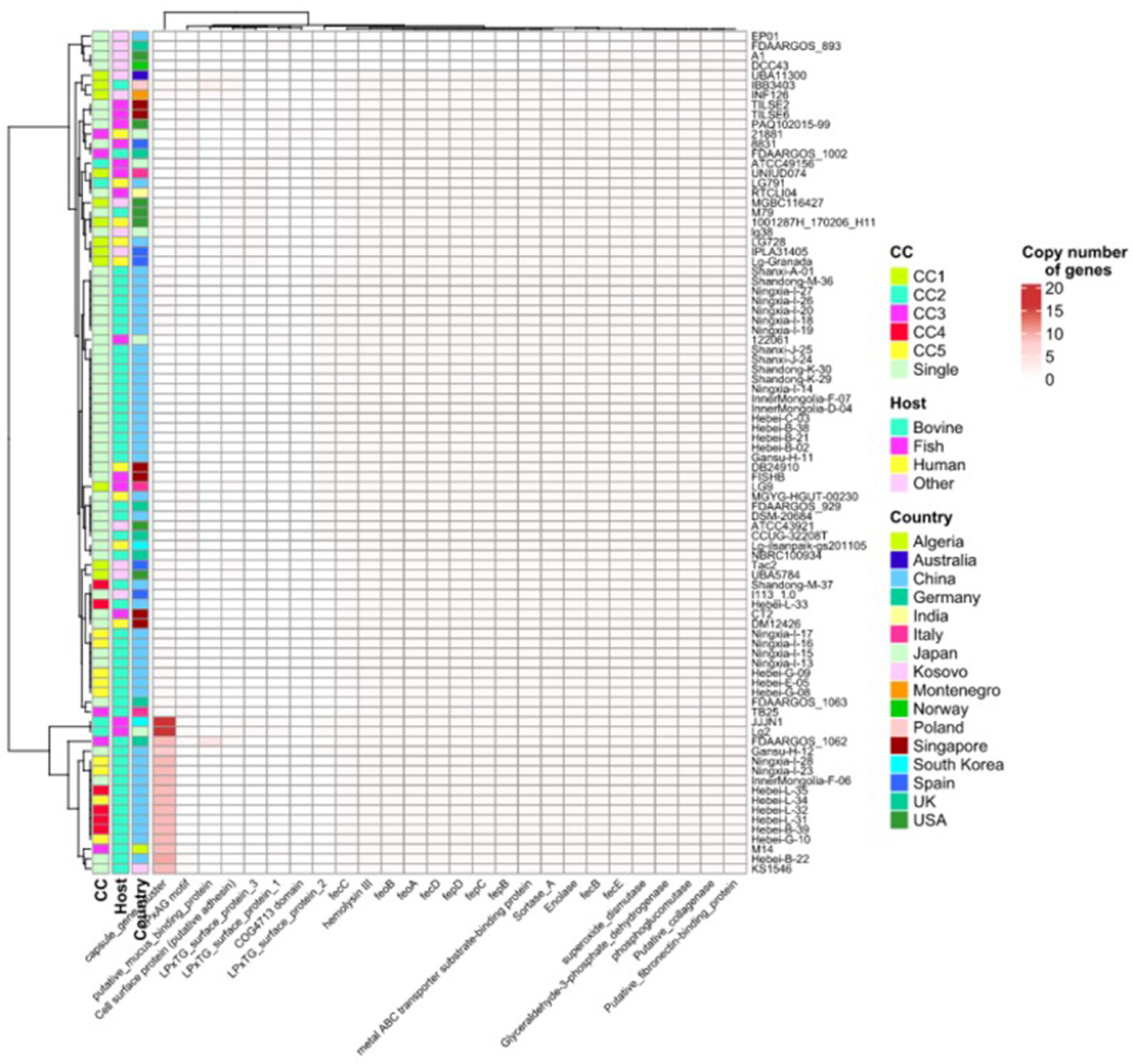
Distribution of virulence factor genes, source of host, country, and clonal complex (CC) of 86 *Lactococcus garvieae* isolates.

Among the 3 toxin-related genes, *hyl-II* was present in all isolates, whereas *hyl-I* and *hyl-III* were absent in ST18 only (represented by 3 isolates: A1, DCC43 and FDAARGOS_893). Nine iron uptake genes (*fepB, fepC, fepD, fecB, fecC, fecD, fecE, feoA,* and *feoB*) were detected in > 81 isolates. In contrast, *fecE* was absent only in EP01. However, *fecB* was absent in EP01 and FDAARGOS_893*. fecD, feoA feoB, fepB, fepC* and *fepD* were absent in 4 isolates: A1, EP01, DCC43 and FDAARGOS_893.

Furthermore*, fecC* was absent in 5 isolates: A1, EP01, DCC43, FDAARGOS_893 plus FDAARGOS_1063. With respect to the 9 adherence-associated virulence factors, *adhPavA* and putative collagenase were identified in all isolates. The gene *adhPsaA* was absent in the same 4 isolates above (A1, EP01, DCC43 and FDAARGOS_893), whereas *Adhesin* was detected in 22 isolates. Genes expressing LPxTG proteins (LPxTG-1, LPxTG-2, LPxTG-3, LPxTG-4, LPxTG-5, and LPxTG-6) were in 4 to 43% of isolates. Six enzyme-related virulence factors were detected in almost all isolates, although *eno* and *srtA* were exclusively absent from 2 (MGBC116427 and UBA11300) and 4 isolates (A1, EP01, DCC43 and FDAARGOS_893). Regarding 16 genes encoding for the capsule gene cluster, the number of gene presented in isolates varied from 0 to 16. Lg2 and JJJN1 were identified with all 16 genes, whereas 47 isolates had 0 genes.

Co-occurrence of virulence genes was visualized in **Fig. 4**, where each box had a phi coefficient value. A phi value of 1.0 indicates a perfect positive relationship between the 2 variables, whereas values **>** 0.7 indicates a fair positive relationship. Most associations among virulence genes were very weak. However, there were strongly positive correlations between 8 genes, including *adhPsaA*, *srtA* and 6 iron uptake genes (*fecD*, *feoA*, *feoB*, *fepB*, *fepC* and *fepD*). In addition, toxin-related genes hyl-I and −3 were also strongly positively associated with each other and those 8 genes. Furthermore, fecC was strongly positively associated with hyl-I, hyl-III and the 8 genes. LPxTG-4 was strongly positively associated with *Adhesin*, whereas *fecB* had strong associations with the 8 genes. However, there were no strong negative associations among the presence of virulence genes.

**FIG 4.**
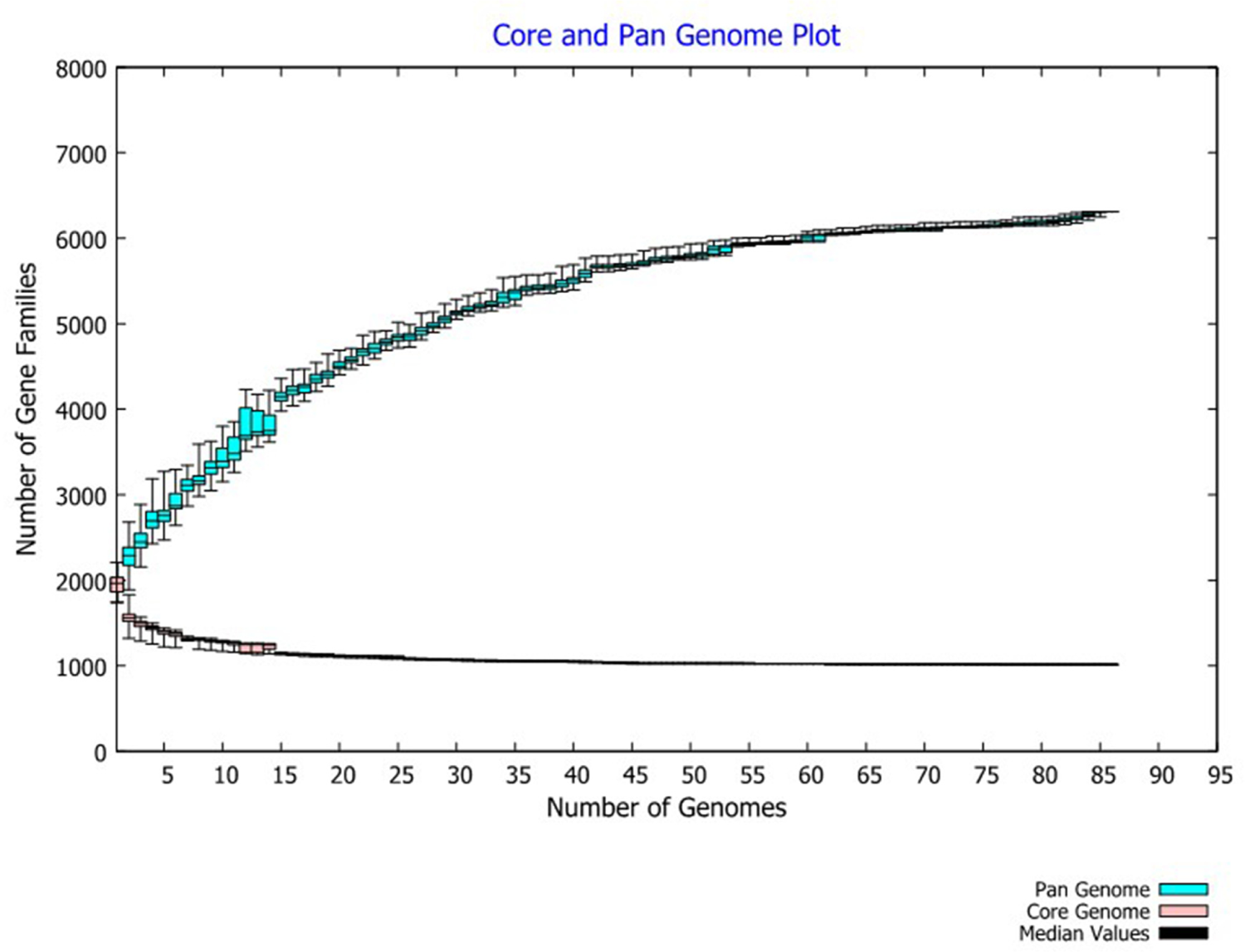
Pan-genome of 86 *Lactococcus garvieae* isolates in this study. The pan-genome consisted of 6310 genes, of which, 1015 core genes, 3641 accessory genes and 1654 unique genes; the size of the genome in the pan-genome increased as the number of isolates increased, but pan-genome size approached convergence. The number of core genes (shared by all isolates) was fairly constant at 1015 genes **(A)**. Distribution of KEGG **(B)** and COG **(C, D)** functional categories in core, accessory and unique genes of 86 *Lactococcus garvieae*.

### Pan-Genome Analyses

The pan-genome of 86 *L. garvieae* isolates tested in this study had 6310 genes. The core genome (shared by 100% isolates) consisted of 1015 genes. The accessory genome (genes in >2 isolates but not in all) consisted of 3641 genes, and the unique genome was composed of 1654 genes. According to BPGA’s calculation, the pan genome was open but approached convergence (**Fig. 4A**).

Functional annotation of genes in the pan-genome performed using the COG and KEGG databases revealed a distribution of functional categories among 3 pan-genome sets (**Fig. 4B, 4C and 4D**). The functions of defense mechanisms, transcription and replication, recombination and repair were enhanced in unique genes, whereas the functions of translation, ribosomal structure and biogenesis were enhanced in core genes in KEGG functional pathways (**Fig. 4B**). The COG functional categories enriched in the unique genome included human disease and membrane transport (**Fig. 4C**). By contrast, COG categories enriched in the core genome included energy metabolism, nucleotide metabolism and translation (**Fig. 4D**).

### Phylogenetic Analyses

A phylogenetic tree was constructed based on the core genes of 86 *L. garvieae* genomes (**Fig. 5**). The longer the branch, the more distant the evolutionary relationship. All trees had 3 clades that contained 4, 38 and 44 isolates respectively. All local isolates except Hebei-B-22 were in the same clades. EP01 was phylogenetically distant from the others. There were signs of host adaptation, which consisted of isolates from various hosts.

**FIG 5.**
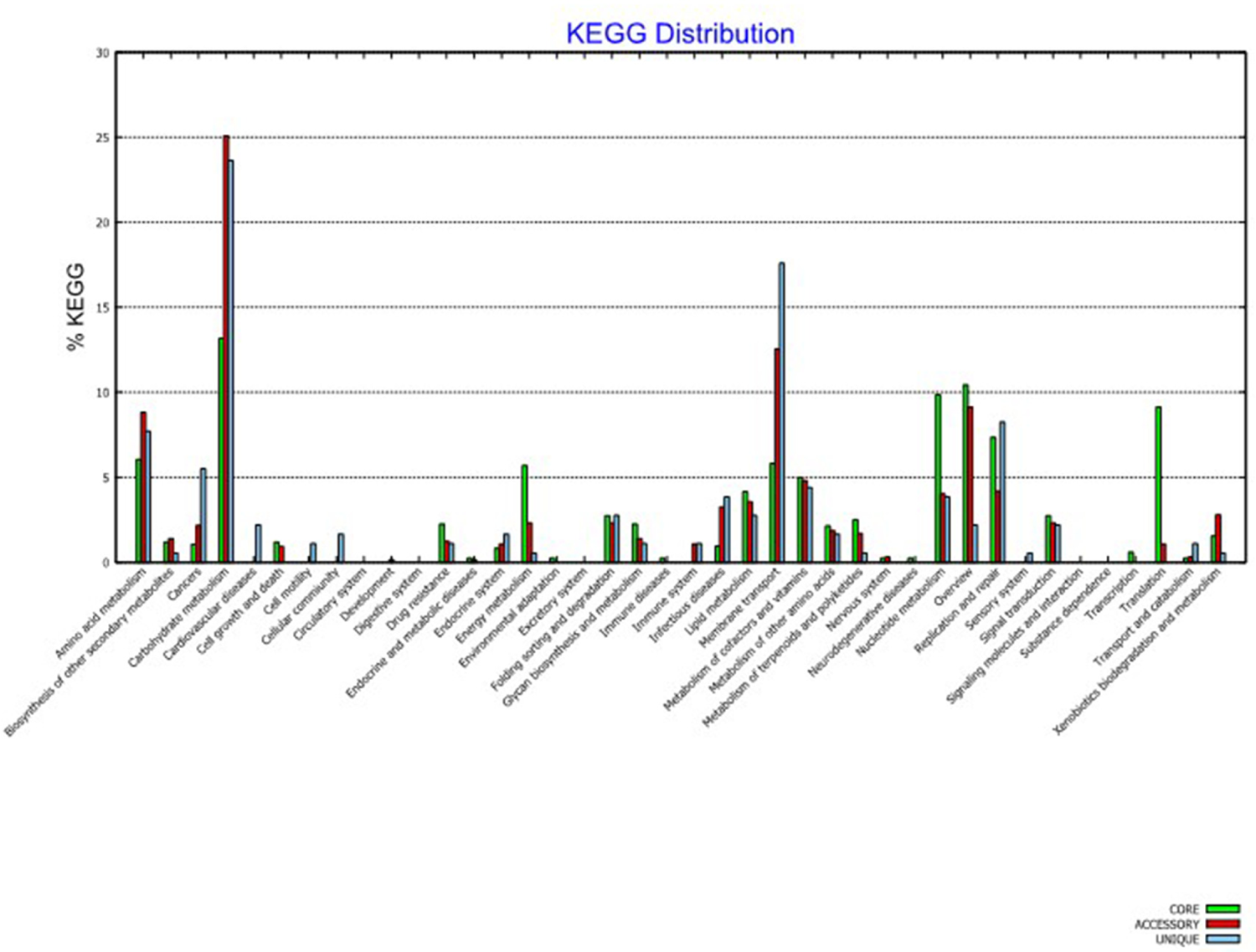
Phylogenetic tree. (A). based on 16S rRNA with source of host (4 hosts, the first ring indicated by a rectangle) and country (16 countries, the second ring indicated by a circle) as well as sequence types (STs, 32 STs, the outermost ring indicated by a triangle) of 86 *Lactococcus garvieae* isolates. **(B).** Phylogenetic tree based on core genes with source of host (4 hosts, the innermost ring indicated by a rectangle) and country (16 countries, the second ring indicated by a circle) as well as sequence types (STs, 32 STs, the outermost ring indicated by a triangle) of 86 *Lactococcus garvieae* isolates.

Phylogenetic tree based on 16s rRNA (**Fig. 5A**) were similar to that using the core genes with minor differences. For example, LG9 was assigned to clade A in core gene based phylogenetic tree but in clade B in 16S rRNA based phylogenetic tree.

Both trees corresponded well with STs and CCs predicted by GrapeTree analysis. For core gene based phylogenetic tree (**Fig. 5B**), all isolates that belonged to the same CC were grouped in the same cluster, except 2 isolates (LG9 and IBB3403) in the 16S rRNA-based phylogenetic tree.

### Pan-Genome-Wide Association Analyses

No significant association was detected between genes and either country or host.

### Core-genome SNPs Analyses

The core-genome single nucleotide polymorphism (SNPs) based phylogenetic tree with metadata annotation is displayed in **Fig. 6**. The numbers of core-genome SNPs among 86 isolates are provided in Supplementary **Table S1**. Several isolates from various hosts were phylogenetically closely related in core SNPs. For example, the number of SNPs between isolates within the same MLST but a different host were: DM12426 (human) and CT2 (fish) was 0, 1001287H_170206_H11 (human) and UBA5784 (metal) was 5, Lg-ilsanpaik-gs201105 (human) and Hebei-B-22 (cow) was 11, 21881 (human) and M14 (cow) was 105, which indicates potential host adaptation in *L. garvieae*.

**FIG 6.**
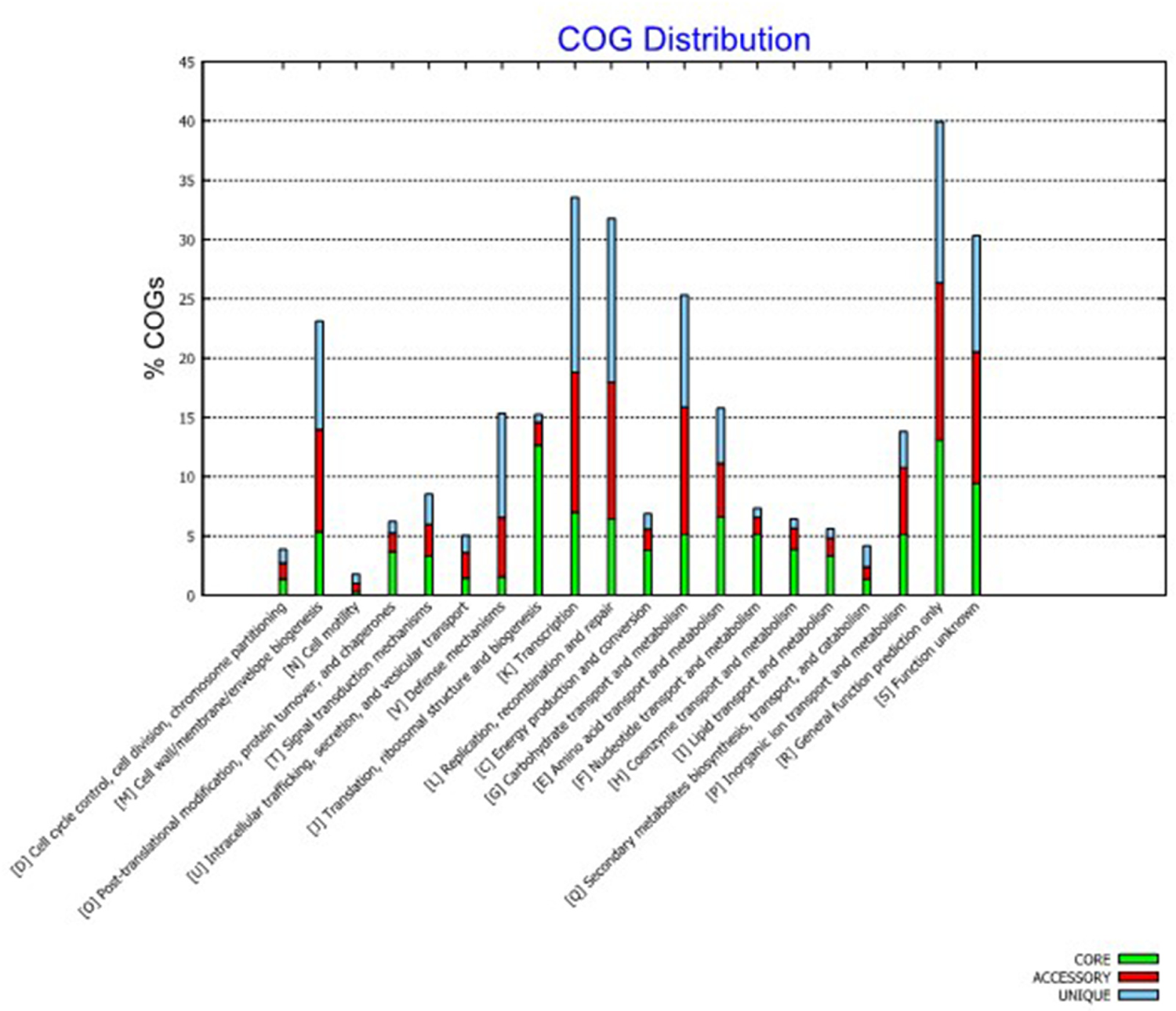
Phylogenetic tree based on core genome single nucleotide polymorphism of 86 *Lactococcus garvieae* isolates.

### Associations between the Co-occurrence of Virulence Genes

Co-occurrence of virulence genes was visualized in **Fig. 7**. SortaseA, LPxTG-6, and adhensin PsaA was in co-occurrence with *fecB*, *fecC*, *fecD*, *feoA*, *feoB*, *fepB*, *fepC*, *fepD*, *hemolysin I*, and *hemolysin III*.

**FIG 7.**
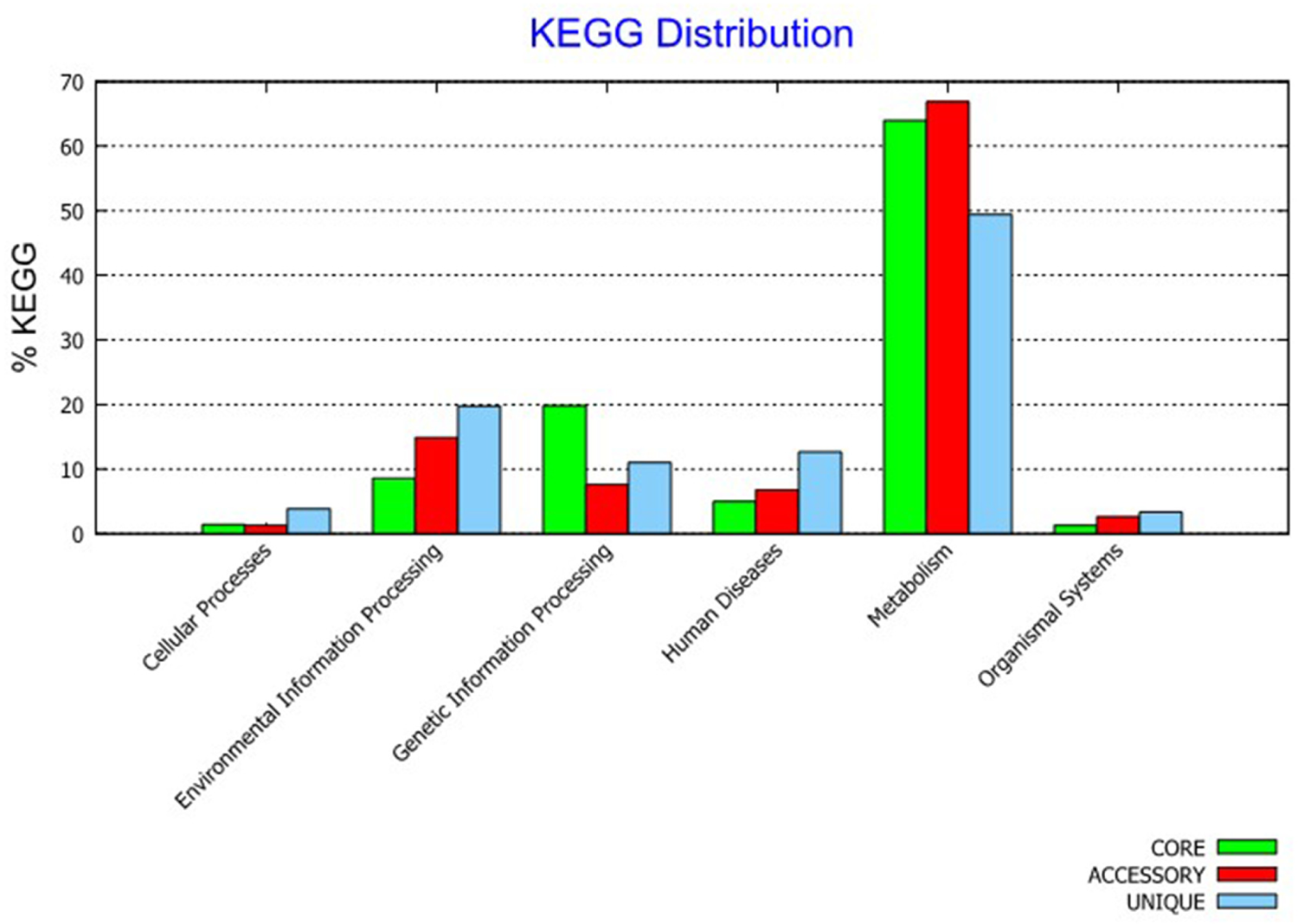
Co-occurrence of virulence genes in 86 *Lactococcus garvieae* isolates.

## DISCUSSION

Although *L. garvieae* was first isolated as a causative agent of bovine mastitis (**27**), most reports have focused more on the epidemiology of fish and human cases. In addition, many studies used sequencing to investigate genotypic characteristics of *L. garvieae* isolates (**8, 28–40**). Therefore, we collected 39 *L. garvieae* isolates from bovine mastitis in China and conducted comparative genome sequence analysis of *L. garvieae*.

The prevalence of *L. garvieae* from clinical mastitis sample was 1.35% during 2017-2021, which increased from 0% in 2017 to 4.10% in 2020 in China. That *L. garvieae* has been misclassified into *Streptococcus spp.* (**41**) has resulted in underreporting of *L. garvieae*. Similarly, the true incidence of human infective endocarditis is difficult to assess due to misidentification with other gram-positive cocci (**42**). This is the first report regarding the prevalence of *L. garvieae* in Chinese dairy herds.

The MLST analysis clustered 86 *L. garvieae* isolates into 32 distinct STs, with 5 CCs and 18 singletons, which is consistent with the population structure in isolates from other hosts (**37, 38, 43**), the environment, or foods (**44**). All strains, except Hebei-B-22, were new STs and phylogenetically close to each other. However, they were distant from isolates of bovine mastitis in other countries, which might indicate geographic effects on the phylogeny. Meanwhile, new STs profiles are comprised of new alleles in gene loci (e.g., *als*, *gyrB* and *galP*), perhaps due to a different evolution rate of those loci (**38**).

Understanding phylogenetic relationships between strains is important for characterizing pathogen transmission. In this study, 3 phylogenetic trees were constructed using core genes and core genome SNPs as well as 16S rRNA, respectively. Core-genes tree and 16S rRNA trees produced similar clades but 16S rRNA failed to resolve relationships toward tree tips. Furthermore, a core-gene tree is in line with MLST and CC. This was not surprising, as 16S rRNA tree is based on only 1 gene, representing only a very small portion of the whole genome. Therefore, many studies recommended core genes for inferring phylogenies (**45, 46**).

Although pathogenicity of *L. garvieae* is poorly understood, some mechanisms have been determined, including presence of a capsule, hemolytic activity via secreted proteins (**47**) and production of siderophores (**48**). Capsulated *L. garvieae* Lg2 was more virulent in fish than the non-capsulated isolate ATCC 49156 (**49**). The capsule gene cluster, located in a genomic island, were identified in Lg2 but absent in ATCC49156, which could be crucial for virulence of *L. garvieae* in fish (**40**). However, existence of the capsule gene cluster has not been detected in all clinical fish isolates from Japan, Spain, Italy, France, Turkey (**22**), USA (**39**), or India (**50**), nor in any human isolates (**31, 69**). In this study, only 2 of 86 isolates had the complete capsule gene cluster, which confirmed that it was not essential for virulence.

Proteases are among the important virulence factors causing rapid and extensive destruction of host tissue. For example, enolase (**50**) can cleave an extracellular proteinaceous matrix and therefore break a host’s structural barrier during colonization. Hemolysin genes might act with secreted proteases to promote host tissue destruction. Genes encoding biosynthesis of iron uptake may be involved in iron acquisition during host colonization (**48**). LPxTG protein (Leu-Pro-any-Thr-Gly), an important virulence factor in *L. garvieae*, binds to the peptidoglycan of cell wall by transpeptidase enzymes called sortases (**40**). In a previous study, *L. garvieae* strain isolated from rainbow trout colonized non-phagocytic cells with the help of LPxTG proteins (**51**). LPxTG proteins and sortases have important roles binding pathogenic bacteria to their host. In this study, genes coding for adhesin, proteases, hemolysin, iron uptake and LPxTG protein were detected in most or all isolates. Strong positive associations within LPxTG-4, *adhPsaA*, *srtA* and iron uptake genes suggest they might act together to promote host tissue destruction and colonization. Gene *pgm,* identified in all isolates, produces protein with an important role in antibody production (**52**).

The minimum inhibitory concentrations (MIC) results were consistent with reports that *L. garvieae* isolates from dairy farms were susceptible to penicillin, ampicillin and amoxicillin-clavulanic acid, imipenem, ceftiofur, enrofloxacin, vancomycin, and marbofloxacin (**10, 53**). However, compared to a previous report (**10**), there were variable degrees of increasing resistant rates for 8 antibiotics, including clindamycin (93.6 to 100%), chloramphenicol (6.4 to 100%), amikacin (2.1 to 90%), cefpodoxime (0 to 82.5%), cephalothin (0 to 10%), cefazolin (40.4 to 45%), gentamicin (0 to 37.5%), and erythromycin (0 to 5%). High resistance of clindamycin has been described as intrinsic for *L. garvieae* and proposed as a selection criterion to distinguish between *L. garvieae* and *L. lactis* (**54**). The *lsaD* gene, identified in most isolates (83/86), could be responsible for intrinsic resistance. *lsaD* is a novel lsa-type family gene detected in lincomycin-resistant strains isolated from fish (**55**). The lsa-type genes are responsible for cross-resistance to lincosamides, streptogramins or pleuromutilins (hereinafter referred to as LSA(P)-resistant phenotype), by coding ATP-binding cassette F proteins in Gram-positive pathogens including Staphylococci, (**56**) Streptococci (**57**), enterococci (**58**), and lactococci (**55**). Increasing resistance against cephalosporins might be related to increasing use of these antibiotics for treatment of infectious diseases on Chinese dairy farms (**59**). The multidrug transporter, *mdtA* is another AMR gene present in most isolates. This gene originally conferred resistance to macrolides, lincosamides, streptogramins and tetracycline in *L. lactis* (**60**), but mutations present in the C-motifs of *mdtA* from *L. garvieae* confer susceptibility to erythromycin and tetracycline (**53**). Furthermore, all 39 isolates with *mdtA* had a limited resistance rate to erythromycin (5%). Some isolates (16/86) harbor the *tetS* gene, including 10 local isolates. Three isolates from the human gut in China, LG729, LG729 and MGYG-HGUT-00230, contained the most abundant AMR genes. Notably, 9 AMR gene were only present in the 3 isolates, including *cat, dfrG, ermA*, *ermB, lunD, fexA, optrA, acc(6’)-aaph(2’’)* and *ant*(*6*)*-la*. The *optrA* gene, first identified in enterococci, has been reported in *Staphylococci*, and *Streptococci*, *Clostridium perfringens* and *Campylobacter coli*; it confers resistance to oxazolidinones and phenicol and has identified on a plasmid of *L. garvieae* (**61**). The spread of antibiotic resistance genes in bacterial populations is aided by various mechanisms of horizontal gene transfer, with plasmid-mediated transfer being the main mechanism for transmission of resistance genes (**62**). Horizontal gene transfer between bacteria is largely mediated by specialized mobile genetic elements, including plasmids, bacteriophages, transposon, insert sequences (IS), intergon, etc., and has been reported in *L. garvieae*. Both *tetS* and *tetM* were associated with conjugative transposon-associated gene in isolates from healthy fish intestines (**63**). Most of the IS in *L. garvieae* had substantial homology to *Lactococcus lactis* elements, implying movement of IS between these 2 species that are phylogenetically closely related (**64, 65**). That these 9 AMR genes were only reported in humans does not support the assertion that AMR genes are transferred to humans from fish or dairy products. Regardless, *L. garvieae* could be a reservoir for antibiotic resistance genes for other bacteria.

In this study, no genes were associated with host specificity, consistent with the phylogenetic analysis and the core-genome SNP analysis that host adaptation occurs in *L. garvieae* isolates. Previous research (**5**) summarized human *L. garvieae* infections associated with consumption of raw fish, seafood, or unpasteurized milk. The core genome SNP analysis underlies the potential host adaptation of *L. garvieae*. Meanwhile, adhesins, haemolysin, fibronectin-binding proteins, penicillin acylase and WxL domain-containing proteins are considered to actively promote bacterial colonization (**66**); most had high similarity across host in those coding sequences. Regardless, underlying mechanisms remain unclear. Consequently, further studies are needed to determine host adaptation mechanisms of *L. garvieae*.

## CONCLUSIONS

This was apparently the first study on comparative genomic analyses of *L. garvieae* isolates from mastitis cows in China. The incidence of *L. garvieae* mastitis was 1.35% in China. Most isolates (38/39) were novel sequence types, 3 antimicrobial resistance genes (*mdtA*, *lsaD* and *tetS)* were identified and there was evidence of host adaptation in these isolates.

## MATERIALS AND METHODS

### Statement of Ethics

This study was conducted in accordance with ethical guidelines and standard biosecurity and institutional safety procedures of China Agricultural University (CAU; Beijing, China). Ethical approval was not needed, as no animal study was involved.

### Sample Collection and Bacteria Identification

Milk samples from clinical cases of mastitis were collected aseptically from dairy cows on Chinese dairy farms and sent to the Mastitis Diagnostic Laboratory at the College of Veterinary Medicine, CAU, Beijing, China. Pathogens were identified by bacteriological culture, colony morphology and 16S rRNA sequencing according to NMC guidelines (**67**). In brief, 50 μl milk was spread on tryptone soy agar with 5% defibrinated sheep blood. The plate was incubated aerobically at 37 °C for 24 h. Bacterial colony morphology was recorded; samples with ≥ 3 morphologically distinct colonies were considered contaminated and excluded from subsequent analyses.

### Antimicrobial Susceptibility Testing

For all 39 *L. garvieae* isolates, MIC of 15 antimicrobials (Chinese National Institutes for Food and Drug Control, Beijing, China), commonly used to treat clinical mastitis in China, were determined by the microbroth dilution method, according to the Clinical and Laboratory Standards Institute (CLSI) guidelines VET01-A4 (CLSI, 2013), with reported breakpoints (**10**). *Staphylococcus aureus* ATCC 29213 was used as the quality control strain.

### Genome Assembly and Annotation

Genomic DNA of putative isolates was extracted using a bacterial DNA extraction kit (TransGen Biotech, Beijing, China) according to the manufacturer’s instruction. Extracted DNA was quantified with a NanoDrop One spectrophotometer (Thermo Fisher Scientific, Waltham, MA, USA) prior to 16S rRNA gene sequencing (Beijing Sunbiotech Inc., Beijing, China). Whole genome DNA was paired-end sequenced (2 × 150 bp) using Illumina NovaSeq 6000 (Illumina, San Diego, CA, USA) at Shanghai Personal Biotechnology Co., Ltd (Shanghai, China). For raw reads, quality control was done with FastQC Version 0.11.9 (https://github.com/s-andrews/FastQC). Low quality bases were trimmed using fastp Version 0.20.1 (https://github.com/OpenGene/fastp) with default settings. Quality trimmed reads were assembled into scaffolds using SPades Version 3.13.1 (https://github.com/ablab/spades) with auto coverage cut-off and shovill Version 1.1.0 (https://github.com/tseemann/shovill) with default settings. Thereafter, 2 assembled scaffolds for each isolate were obtained, and a draft genome of each isolate was selected using Quast Version 5.0.2 (https://github.com/ablab/quast) with N50, L50 from the above-mentioned 2 assembled scaffolds. Assembly completeness was assessed using Busco Version 5.2.2 (https://github.com/WenchaoLin/BUSCO-Mod) with reference to lineage lactobacillales_odb10. Only genomes with completion ≥ 95% were considered “high-quality draft genome” and were included in further analyses (**68**). In addition, whole genome sequence assemblies fasta files of 51 *L. garvieae* (accessed on March 24, 2022) were downloaded from NCBI. To ensure high-quality genomes, all genomes were analyzed by BusCom Version 5.2.2 (lineage lactobacillales_odb10) and 3 assembles were excluded from subsequent analysis. There were 3 ATCC 49156 assemblies; we chose the 1 with the highest assembly level. Therefore, a total of 39 isolates from composite (or quarter) milk samples and 47 assemblies from NCBI were obtained in the subsequent genome annotation and pan-genome analysis. Annotation of the genome was performed using Prokka Version 1.14.6 (https://github.com/tseemann/prokka) with default settings. MLST analyses Multilocus sequence typing (MLST) using whole genome sequences was performed to determine sequence types (ST) of the 86 isolates. A *L. garvieae* MLST database was constructed based on reported datasets (**69**) as there was no publicly available MLST scheme for *L. garvieae*. Similarly, the database was integrated into ABRicate local database, and we aligned the 86 *L. garvieae* genomes against the dataset by ABRicate. Sequence types were assigned to new allele patterns and added to the existing MLST scheme for *L. garvieae* constructed by (**69**). Clonal complex (CC) was defined as a group of STs in which every ST shared at least 5 of 7 identical allele profiles with at least 1 other ST in the group. The minimum spanning tree (MST) was constructed by the goeBURST algorithm and visualized with the PhyloViz web server (https://online.phyloviz.net/index) to infer phylogenetic relationships among STs.

### Identification of Antimicrobial Resistance Genes and Virulence Factors

Antimicrobial resistance genes were identified by blasting *L. garvieae* genomes against ResFinder database via Resfinder Version 4.1.11 (https://cge.cbs.dtu.dk/services/ResFinder/) and The Comprehensive Antibiotic Resistance Database (https://card.mcmaster.ca/) via RGI Version 5.2.1 (https://github.com/arpcard/rgi). A set of virulence genes of *L. garvieae* was summarized from previous reports (**22, 69, 70**); they included haemolysin I, II and III (*hlyI*,-*II* and *-III*), iron uptake genes (*fepB*, *fepC*, *fepD*, *fecB*, *fecC*, *fecD*, *fecE*, *feoA*, and *feoB*), capsule gene cluster (CGC), adhesions (*adh*, *adhPavA*, *adhPsaA*), putative collagenase (*colA*), LPxTG surface proteins 1, 2, 3, 4, 5, 6 (LPxTG-1, LPxTG-2, LPxTG-3, LPxTG-4, LPxTG-5 and LPxTG-6) and enzyme-related virulence factors NADH oxidase(*Nox*), glyceraldehyde-3-phosphate dehydrogenase (GAPDH), phosphoglucomutase (*Pgm*), superoxide dismutase (*Sod*), enolase (*Eno*) and SortaseA (*srtA*). The database was integrated into the ABRicate local database. We blasted the 86 (39 from our study and 47 from NCBI) *L. garvieae* genomes against the database using ABRicate to determine virulence genes. The presence of antimicrobial resistance or virulence gene was defined using the cut-off value of 80% sequence coverage and 80% nucleotide identity (ABRicate default settings).

### Pan-genome Analyses

The pan-genome of 86 *L. garvieae* isolates was computed using BPGA Version 1.3 (https://iicb.res.in/bpga/) with USEARCH algorithm to cluster orthologous gene families using faa files of local isolates produced by Prokka and retrieved from NCBI directly. For BPGA analysis, a core gene was defined as a gene present in all the genomes; an accessory gene was present in > 1 genome but not all genomes; and a unique gene was only present in a single genome. Functional annotations of core, accessory, and unique genes were obtained after comparing sequences to COG and KEGG databases incorporated in BPGA Version 1.3.

### Phylogenetic Analyses

A 16S rRNA phylogenetic tree was constructed based on 16s rRNA genes. In addition, we also constructed another phylogenetic tree using alignment of core genes produced by BPGA. For 16S rRNA phylogenetic analysis, Barrnap Version 0.9 (https://github.com/tseemann/barrnap) was used to extract 16S rRNA gene from the whole genome sequence. The 16S rRNA gene sequences were edited and aligned using MAFFT multiple sequence alignment algorithm (stargety “L-IINS-I”; https://github.com/GSLBiotech/mafft). Maximum-likelihood (ML) trees based on this alignment were constructed using FastTree Version 2.1 (https://github.com/PavelTorgashov/FastTree). Visualization of the phylogenetic tree was performed using iTOL (https://itol.embl.de/) with metadata (clonal complex, source of host and country) of the isolates.

### Pan-genome Wide Association Analyses

To identify genes potentially associated with traits, such as host, clonal complex, and country, we performed pan-genome wide association analysis using Scoary. Annotation of whole genome sequences of the 86 *L. garvieae* isolates were performed using Prokka Version 1.14.6 (https://github.com/tseemann/prokka), and the resultant gff files were used for pan-genome analysis with Roary Version 3.13.0 (http://sanger-pathogens.github.io/Roary/) to produce gene presence and absence data. Thereafter, genes associated with host, country, ST or clonal complex were identified with Scoary VVersion 1.6.16 (https://github.com/AdmiralenOla/Scoary). Categorical traits were dichotomized prior to pan-genome wide association analysis with Scoary.

### Core-genome SNPs Analyses

In addition to pan-genome wide association analyses, we also performed core-genome SNPs analyses. Core genome alignment and single nucleotide polymorphism (SNP) were detected for all 86 genome sequences using parsnp Version 1.7.2 (https://github.com/marbl/parsnp). Meanwhile, the exact numbers of SNPs among genomes from various hosts in the same MLST group and closely related in core-gene based phylogenetic tree were determined with snp-dists Version 0.8.2 (http://sanger-pathogens.github.io/snp-sites/) using the sequence alignment file produced from parsnp. Phylogenetic tree based on core genome SNPs was annotated with iTOL.

### Associations between the Co-occurrence of Virulence Genes

Co-occurrence of virulence genes was determined with phi coefficient using the Phi function in psych package Version 2.2.5 (https://cran.r-project.org/web/packages/psych/) with R Version 4.1.3 (https://www.r-project.org/) and *P* < 0.05 was considered significant in a 2-tailed test. The pair-wised phi coefficients between the presences of virulence genes were visualized using ggplot2 Version 3.3.6 (https://cran.r-project.org/web/packages/ggplot2/).

### Data Availability

All whole genome sequence data used in this study are available without restriction from NCBI under BioProject no. PRJNA848370.

## ACKNOWLEDGMENTS

This study was supported financially by the following: National Natural Science Foundation of China (No. 32172928), the High-End Foreign Experts Recruitment Program (No. 110000202720220022), and Beijing Municipal Natural Science Foundation (No. 6222031). B.H. and Y.L. designed and supervised the study. Y.L., J.H., Y.W., J.G. and S.Q. performed the experiments, and wrote the manuscript. Z.D. and H.W.B. assisted in the analyses and re-edited the manuscript. B.H., J.P. K. and H.W.B. revised the manuscript. All authors read and approved the final manuscript. The authors declare that the research was conducted in the absence of any commercial or financial relationships that could be construed as a potential conflict of interest.

## SUPPLEMENTAL MATERIAL

Supplemental material is available online only. **Table S1.xlsx** file, 45 KB.

---

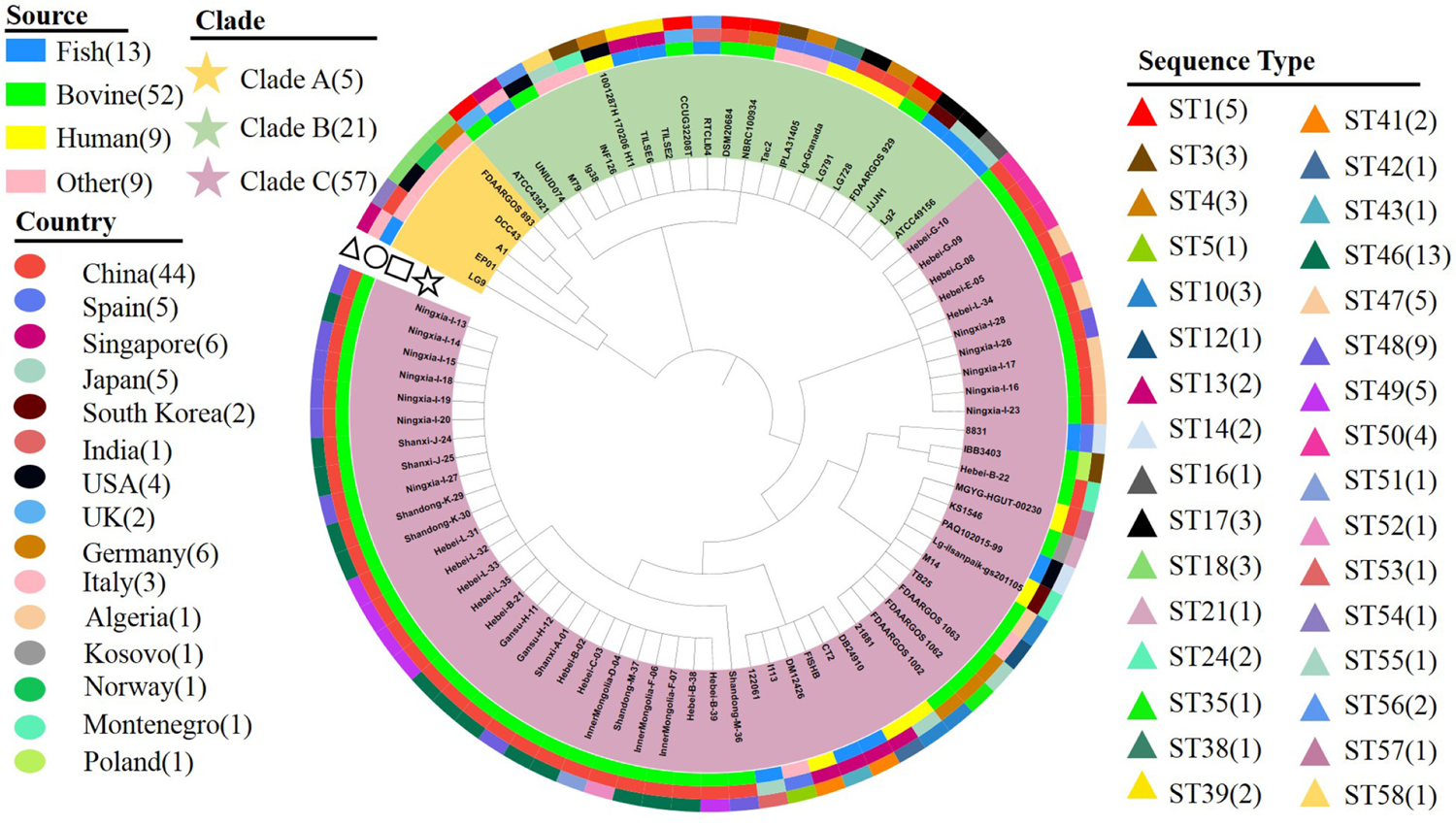

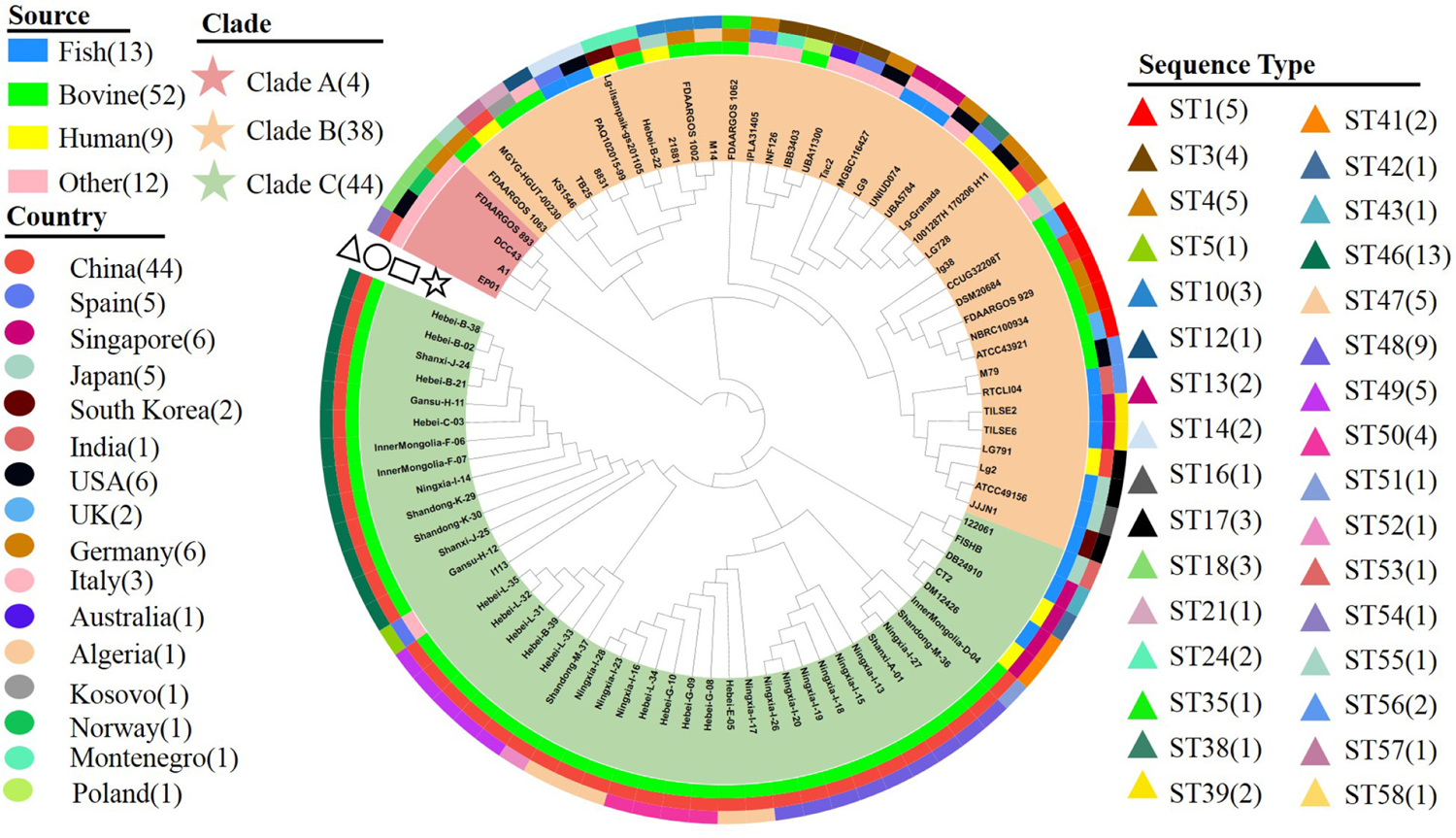

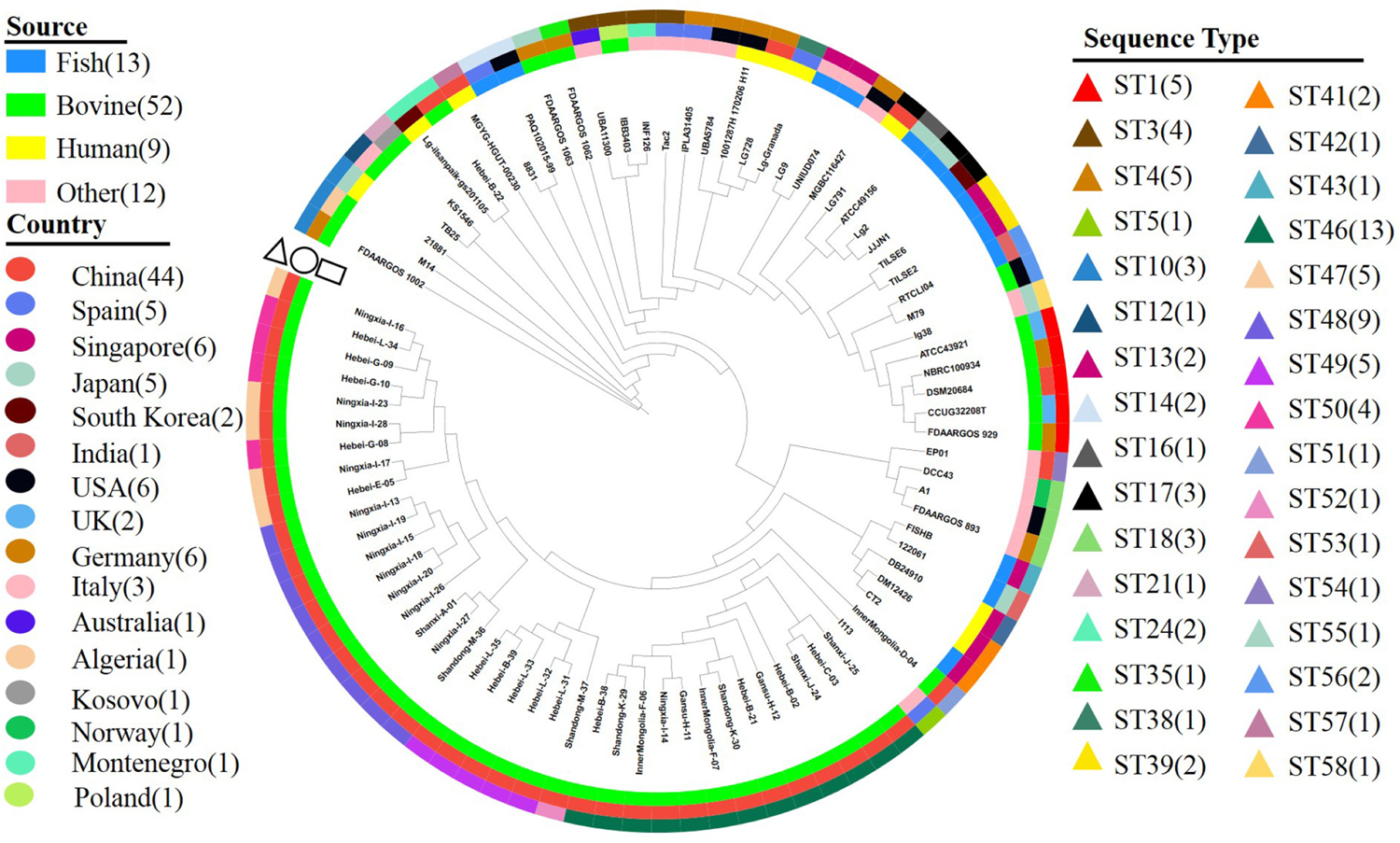

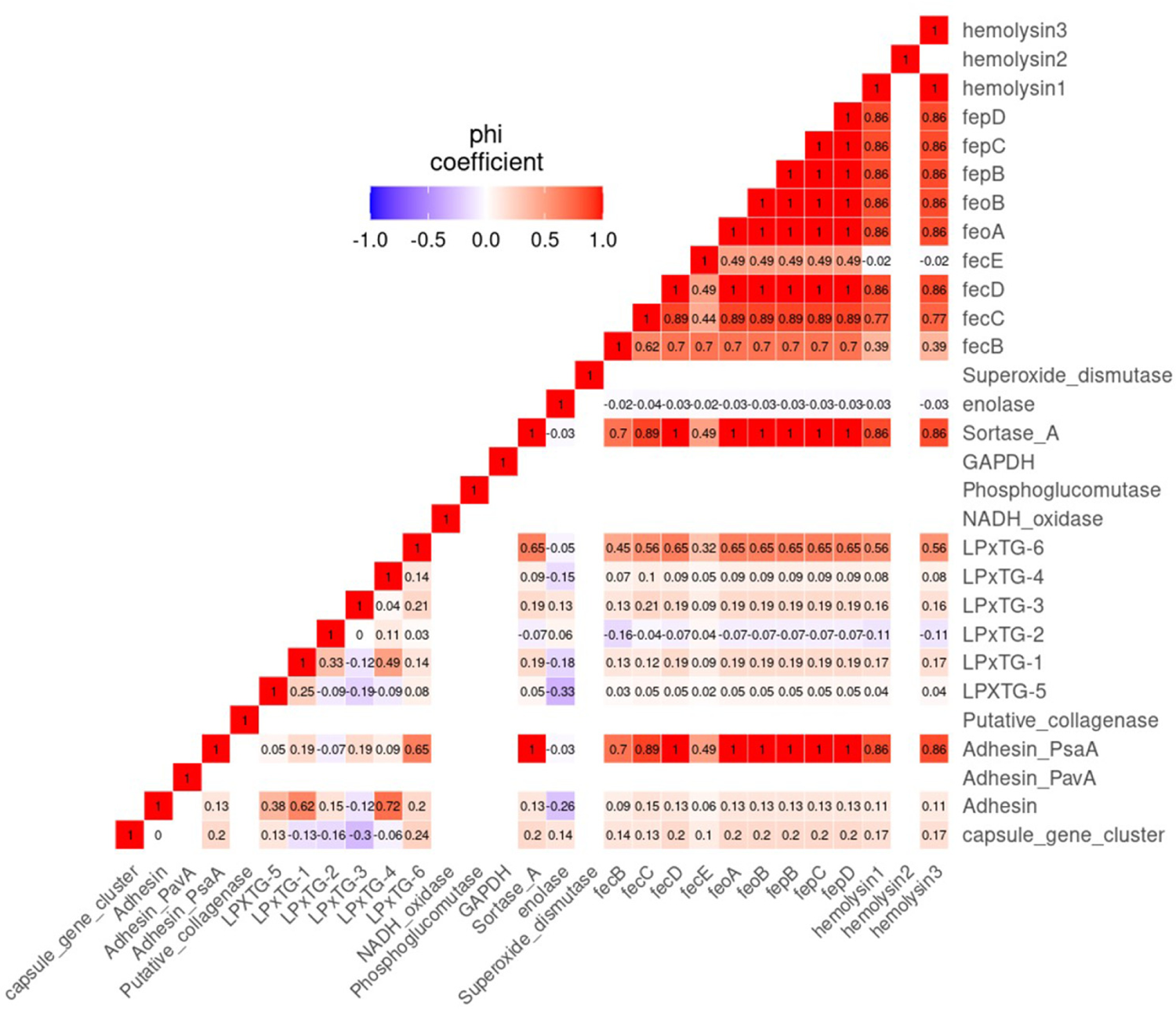

